# Critical role of growth medium for detecting drug interactions in Gram-negative bacteria that model *in vivo* responses

**DOI:** 10.1101/2022.09.20.508761

**Authors:** Kathleen P. Davis, Yoelkys Morales, Anne L. McCabe, Joan Mecsas, Bree B. Aldridge

## Abstract

The rise in infections caused by multidrug-resistant (MDR) bacteria has necessitated a variety of clinical approaches, including the use of antibiotic combinations. Antibiotic susceptibility is affected in part by the growth state of bacteria within various tissues. Here we tested the hypothesis that drug-drug interactions vary in different media, and hence, using a medium that reflects tissue environments will better predict *in vivo* outcomes. We systematically studied pair-wise antibiotic interactions in three different media (CAMHB, a urine mimetic, and a lung mimetic) using three Gram-negative ESKAPE pathogens, *Acinetobacter baumannii* (Ab), *Klebsiella pneumoniae* (Kp), and *Pseudomonas aeruginosa* (Pa). There were pronounced differences in responses to antibiotic combinations between the three bacterial species grown in the same medium. However, within species, Pa responded to drug combinations similarly when grown in all three different media, whereas Ab responded similarly when grown in CAMHB and a lung mimetic medium. By contrast, drug interactions in Kp were poorly correlated across three different media. To assess whether distinct media were predictive of antibiotic interactions in Kp in the lungs of mice, we developed a treatment strategy and tested three antibiotic combination pairs. Measurements obtained *in vitro* from lung mimetic medium, but not rich medium, predicted *in vivo* outcomes. This work demonstrates that antibiotic interactions are highly variable when comparing across three gram-negative pathogens and highlights the importance of growth medium by showing a superior correlation between *in vitro* interactions in a growth medium that resembles the tissue environment and *in vivo* outcomes.

## Main Text

### Introduction

The rise in antimicrobial resistant (AMR) infections is a global health crisis that threatens the ability to treat many bacterial, viral, and fungal infections (1). The rate of multi-drug resistant (MDR) bacterial infections has been steadily increasing for the past 20 years and was boosted by the recent SARS-CoV2 pandemic, which saw a significant increase in MDR related secondary bacterial infections leading to increased rates of morbidity and mortality (2). Among MDR infections, some of the most harmful and difficult to treat are those caused by Gram-negative bacteria within the category of ESKAPE pathogens (*Enterococcus faecium*, *Staphylococcus aureus*, *Klebsiella pneumoniae* (Kp), *Acinetobacter baumannii* (Ab), *Pseudomonas aeruginosa* (Pa), and *Enterobacter* species). This group of bacteria is recognized by the World Health Organization (WHO) as capable of pan-resistance (3). Clinicians and scientists have responded by investing in antibiotic stewardship and novel therapies to combat these infections (4).

Treatment of infections caused by MDR pathogens often involves the use of a combination of antibiotics to limit the emergence of antibiotic resistance (5). Though current meta-analyses and clinical trials sometimes support the use of combination therapy for Gram-negative MDR infections, these studies are often inconclusive (6). Limitations to our understanding of drug combination therapies for the treatment of MDR infections inhibits our ability to effectively predict *in vivo* efficacy using *in vitro* assays (7). Therefore, therapy is reliant on clinical reasoning by individual physicians on a case-by-case basis. Another barrier for combination therapy testing is the resources required to test combinations of antimicrobial in a traditional plate-based checkerboard assay due to the exponential cost of testing multiple drugs in high-order combinations (8). Understanding the potential for combination therapy is further complicated by the fact that these Gram-negative bacteria can cause infections at multiple sites (3). Thus, standard rich media conditions (as defined by the European Committee on Antimicrobial Susceptibility Testing and the International Organization for Standardization) used for *in vitro* assays may not effectively predict how a combination will interact in various distinct *in vivo* environments where bacterial metabolism can differ (9). Here, we hypothesize that environmental effects on bacterial physiology influence drug interactions and that measurements taken using tissue mimetic media will be better able to predict *in vivo* outcomes.

To test this hypothesis, we undertook a systematic study of pairwise antibiotic interactions in different growth environments focused on three Gram-negative ESKAPE pathogens, Ab, Kp, and Pa, which together are among the major worldwide causes of nosocomial infections (3). We measured synergistic, additive, and antagonistic antibiotic interactions in standard rich medium and compared these to measurements made in media designed to simulate lung or urine environments, to model two common sites of infection for these three pathogens. Generating this large dataset of antibiotic interaction measurements allowed us to interrogate the combinatorial space for these species through several lenses. We first asked whether antibiotic combinations behave similarly across all species and media conditions. After finding only one instance where the outcome of an antibiotic combination was similar for all three species grown in all three media, we next teased out both media-specific and species-specific contributions to this observation. Comparisons between different species grown within the same media conditions generally showed poor correlations. However, we did observe reasonable correlations for Pa between the three different media conditions, and a strong correlation between Ab responses in CAMHB and the lung-like condition. By contrast, Kp had very poor correlations across all three conditions. We then assessed the capacity to translate *in vitro* measurements made using medium predicted to simulate the lung nutritional environment or standard rich medium to results found in mouse lung infections. For Kp, *in vitro* measurements in a lung mimetic medium were significantly more predictive of *in vivo* results. This work demonstrates that antibiotic interactions are highly variable when comparing across three gram-negative ESKAPE pathogens and highlights the importance of growth medium by showing a superior correlation between *in vivo* interactions and *in vitro* interactions in a tissue mimetic growth medium.

## Results

### Systematic survey of condition-specific drug interactions in three Gram-negative pathogens

To determine the dependence of drug interactions on growth conditions and bacterial species, we generated a dataset of pairwise drug interaction measurements from a panel of drugs that were tested against Ab, Pa, and Kp (Figure 1A). We chose well-characterized strains of each species – Ab strain ATCC 17978, Pa strain PaO1, and Kp strain ATCC 43816. Each of these strains was grown in three different growth conditions (Fig. 1A) and tested against clinically relevant drugs that cover a range of classes and mechanisms of action (Table 1) (10). The drugs trimethoprim-sulfamethoxazole (BAC) and cefixime were tested only with Kp because of their clinical relevance specific to Kp (11). We systematically measured drug interactions among strains and media conditions by leveraging the efficiency of a methodology called <u>dia</u>gonal <u>m</u>easurements <u>o</u>f <u>n</u>-way <u>d</u>rug interactions (DiaMOND), which implements a geometric optimization of the standard checkerboard assay (12, 13).

**Table 1:** Antibiotics used in this study.

| **Antibiotic** | **Abbreviation** | **Class** | **Mechanism of Action** |
| --- | --- | --- | --- |
| trimethoprim-sulfamethoxazole | BAC | antifolate antibacterial (trimethoprim); sulfonamide (sulfamethoxazole) | folate synthesis inhibition |
| cefepime | CEF | cephalosporin | cell wall synthesis inhibition |
| cefixime | CFX | cephalosporin | cell wall synthesis inhibition |
| colistin | COL | Polymyxin | cell membrane disruption |
| ceftriaxone | CTX | cephalosporin | cell wall synthesis inhibition |
| gentamicin | GEN | aminoglycoside | protein synthesis inhibition |
| levofloxacin | LEV | fluoroquinolone | inhibition of DNA replication and transcription |
| meropenem | MER | Carbapenem | cell wall synthesis inhibition |
| rifampicin | RIF | antimycobacterial | RNA synthesis inhibition |
| tigecycline | TIG | glycylcycline | protein synthesis inhibition |

Class, mechanism of action and 3-letter abbreviation of all antibiotics used in this study.

**Figure 1:**
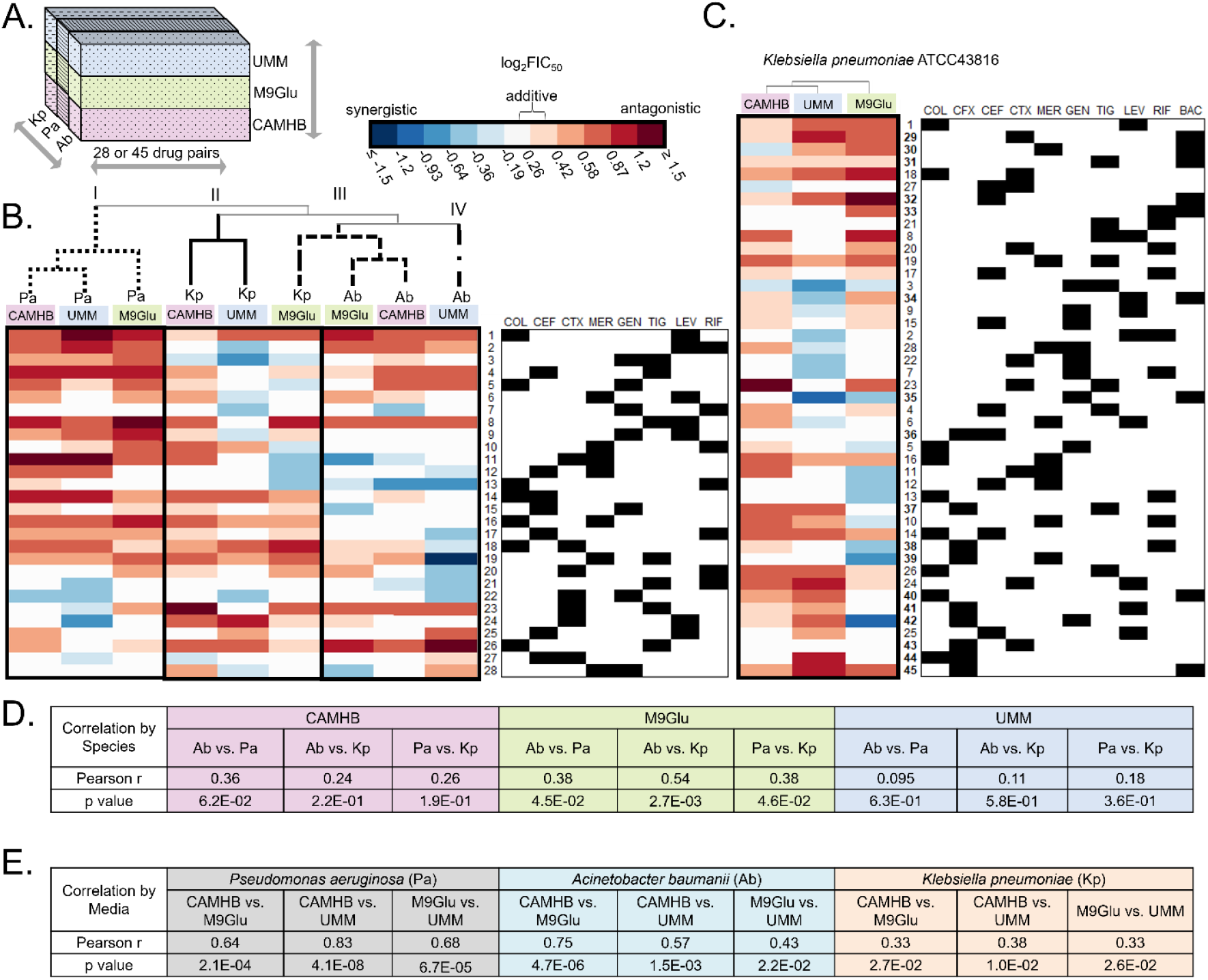
Variation in drug interaction across species and media. (A) Study design involved testing 28 pairs of antibiotics against *A. baumannii* ATCC17978 (Ab) and *P. aeruginosa* PaO1 (Pa), and 45 pairs against *K. pneumoniae* ATCC 43816 (Kp). Testing was done with all strains grown in three different growth conditions: cation-adjusted Mueller-Hinton Broth (CAMHB); M9 minimal medium + 0.5% glucose supplemented with 0.6μM iron (II) sulfate, and with 10mM sodium acetate for Ab and Pa; and urine mimetic medium (UMM) (Brooks & Keevil, 1997) supplemented with 0.01% glucose and 0.6μM iron (II) sulfate for Kp. (B) Clustergram of the log_2_FIC_50_ values of the 28 drug pairs tested across all three species and media, with each row representing a drug combination (indicated by black squares under the drug abbreviations in the table on the right) and each column representing a species tested in a particular medium. Each drug-pair number is maintained throughout the manuscript for ease of comparison. Hierarchical clustering was performed using average linkage between clusters and Pearson correlation distance metric between columns. Each value represents an average of at least three replicates (C) Clustergram of the log_2_FIC_50_ values of all 45 drug pairs tested against Kp in all three media (columns). Hierarchical clustering, notation of drug pairs, and representation of log_2_FIC_50_ is the same as for (B). (D) The Pearson correlation coefficient (r) and p value were determined for each species-to-species comparison of mean log_2_FIC_50_ values in the three media conditions. (E) The Pearson correlation coefficient (r) and p value was determined for each medium-to-medium comparison of mean log_2_FIC_50_ values in the three species.

Each of these species can cause infection at multiple sites in the body, which have different growth conditions that may influence bacterial metabolism (14, 15) and drug response (16–18). However, to our knowledge the effect of growth conditions on drug interactions across different species has not been tested systematically. To directly evaluate whether different growth conditions impact pairwise drug interactions, we employed three media conditions – Cation-Adjusted Mueller Hinton Broth (CAMHB), M9 + 0.5% Glucose + Fe(II)SO4, pH 7.0 (M9Glu), and Urine Mimetic Media (UMM), which has a pH of 6.4 with creatinine and urea as the predominate carbon sources (see Methods). We chose CAMHB because it is a standard for microbiological susceptibility testing, and it is a rich medium that is high in amino acids and vitamins (19). M9 supplemented with 0.5% glucose and 0.6μM Fe(II)SO4 is a minimal medium that lacks amino acids yet still produces consistent reproducible bacterial growth. Additionally, M9Glu reflects the low amino acid availability observed in bronchoalveolar lavage fluid from mice (20, 21) making it a better mimic for bacterial infection in the lungs and other amino acid deficient environments. Finally, we used a Urine Mimetic Media based Brooks & Keevil (1997) to approximate the growth environment experienced by the bacteria during a urinary tract infection (22).

We generated a drug interaction dataset using DiaMOND (12, 23) by measuring the three most information-rich dose response curves: the combination dose responses of increasing equipotent doses of two drugs, and the dose responses of each single drug. We use these dose response curves to calculate the fractional inhibitory concentration (FIC), a measure of drug interactions. The FIC is the ratio of the observed combination dose that results in a certain level of growth inhibition compared to the expected combination dose if the two drugs are additive (see Materials and Methods). Here, we report log transformed FIC scores; log_2_FIC_50_ scores close to zero indicate additivity, more negative scores indicate synergy (e.g., the drugs combined are more effective than expected based on their efficacies alone), and more positive scores indicate antagonism between drug pairs. The efficiency of DiaMOND enabled us to create a dataset of >300 unique combinations of species, medium, and pairwise drug interactions.

### Drug interactions are dependent on species and growth environment

The drug interaction data for 28 drug pairs tested for three species, each grown in three media conditions, is shown in a heatmap with hierarchical clustering in Fig. 1B (log_2_FIC_50_). We observed that for each medium (color-coded above the clustergram), drug combinations varied in their log_2_FIC_50_ scores among different species. Furthermore, within individual species, drug combinations often varied in their log_2_FIC_50_ scores across the three media conditions (three columns within a black box), although the extent of this variation is different for different species. The three Pa growth conditions cluster together (Cluster I), indicating their similarity to each other and differences from Ab and Kp. On the other hand, Ab CAMHB and M9Glu cluster together with Kp M9glu (Cluster III), while Ab UMM is in an adjacent cluster (Cluster IV). Kp CAMHB and Kp UMM make up Cluster II. These clustering patterns suggest that drug interactions are influenced by differences between species while the impact of media is more pronounced in some species versus others.

Pearson correlation coefficients were derived to quantify changes in drug interactions between different species in the same growth conditions (Fig 1D) and between different growth conditions within each species (Fig 1E). The outcome of drug pair interactions between species within the same medium was extremely variable; all nine Pearson coefficients were below 0.6 and eight of the nine were below 0.4 (Fig. 1D). Thus, species-specific attributes impact drug interactions under these conditions. Curiously, despite the overall low Pearson coefficients, the correlation between species was consistently highest in M9Glu and lowest in UMM (Fig. 1D). In contrast to differences in drug interactions between species within the same medium, drug interactions in Pa between all three media showed high and significant correlations, with all three correlations above 0.64 (Fig. 1E). This was also reflected visually by the clustergrams (Fig. 1B). Likewise, drug interactions in Ab between CAMHB and M9Glu showed a high and significant correlation (Fig. 1E). On the other hand, Kp had low correlations in medium-to-medium comparisons, with all three correlations below 0.4 (Fig. 1E). In summary, drug interactions varied widely across different species, while media conditions had larger effects on drug interactions in some species compared to others.

### Drug interactions are overall biased towards antagonism, but synergy is more prevalent in some species in nutrient-depleted media

Efforts to develop clinically impactful combination therapies are focused on identifying synergistic combinations. Though we did not find combinations that were synergistic across all species and conditions tested, one combination, ceftriaxone + gentamicin (#22), was synergistic in a single medium (UMM) across all three species. Additionally, two combinations, colistin + rifampicin (#13) in Ab and gentamicin + tigecycline (#3), were synergistic across all three growth conditions for Kp. However, combinations that were synergistic against one species in a particular growth condition were often not synergistic against other species in that growth condition nor in a different growth condition for the same species (e.g., meropenem plus tigecycline (#19) was synergistic for Ab in UMM, but antagonistic for Ab in the other growth conditions, and antagonistic for Pa and Kp in UMM). One combination was antagonistic across all species and media, colistin + levofloxacin (#1). The tendency towards antagonism was dependent on growth conditions, with combinations in UMM less likely to be antagonistic than those in CAMHB or M9Glu. Specifically, nine combinations were antagonistic across all three species in CAMHB (#’s 1, 4, 5, 6, 8, 14, 18,19, and 26), eight in M9Glu (#’s 1, 4, 8,18, 19, 20, 23, and 26), and one in UMM (#1) (Fig. 1B).

Despite the overall predominance of antagonism, the number of combinations that were additive or antagonistic in CAMHB but synergistic in one or both nutrient-depleted media differed for the three species (Fig. 2, gray regions of graphs). For Pa, four combinations that were additive in CAMHB were synergistic in UMM (Fig. 2B). For Ab, more combinations shifted from additive or antagonistic in CAMHB to synergistic in a nutrient-depleted media: two synergies were identified in M9Glu (Fig. 2C gray region) and six synergies were found in UMM (Fig. 2D gray region). For Kp, among the drug pairs tested in all three species, five combinations were synergistic in M9Glu but not in CAMHB (Fig. 2E gray region), and six combinations were synergistic in UMM but not in CAMHB (Fig. 2F gray region). Among the drug pairs tested only in Kp (the 8 core drugs against trimethoprim-sulfamethoxazole or cefixime), four were synergistic in M9Glu but not CAMHB, and two were synergistic in UMM but not CAMHB. Thus, for Ab and Kp tissue mimetic conditions revealed additional synergistic combinations and may yield different predictions than measurements made in CAMHB. On the other hand, it may be sufficient to test drug pairs in rich media alone for Pa.

**Figure 2:**
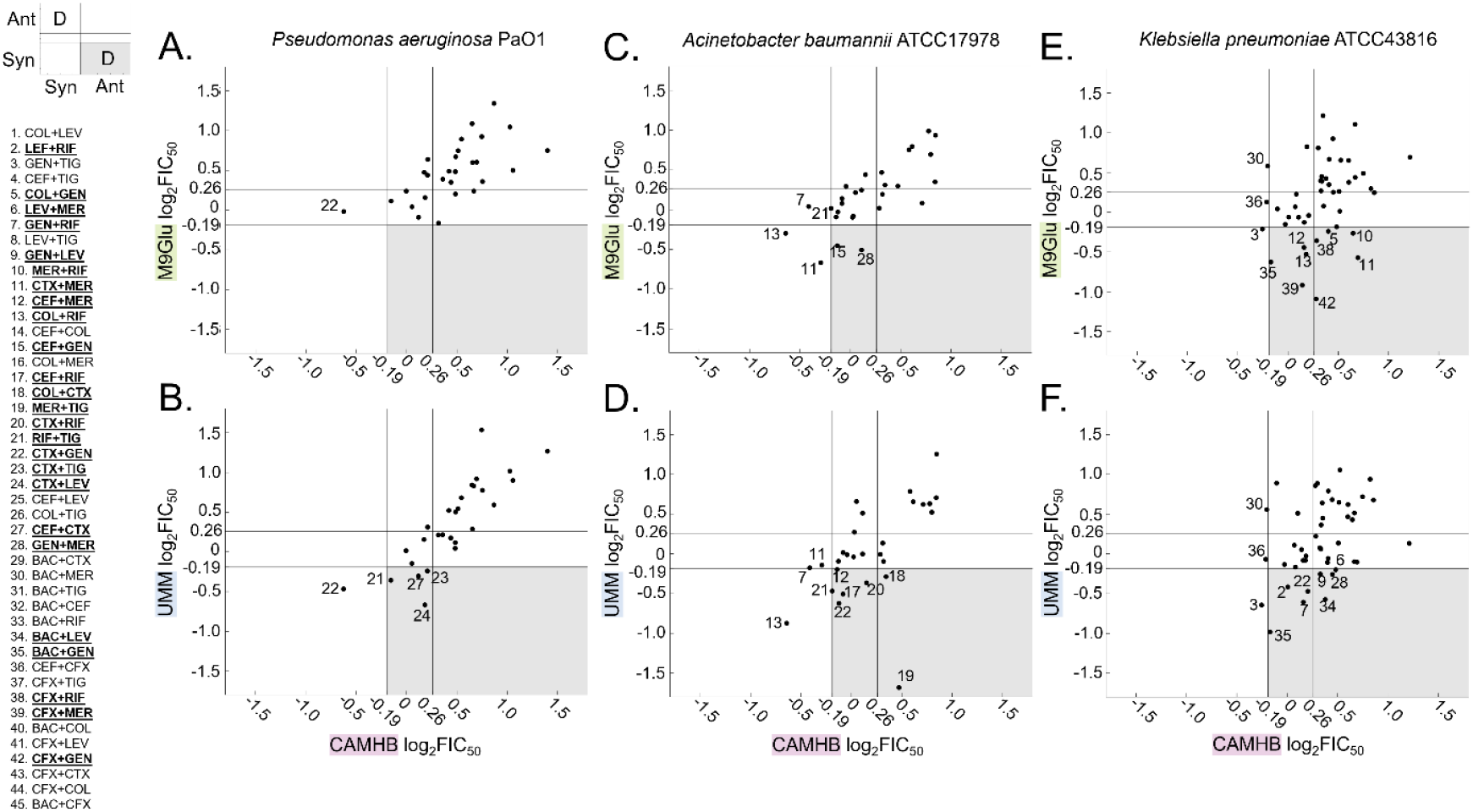
Nutrient-depleted media (M9Glu, UMM) reveal more synergistic combinations than standard rich media (CAMHB) for some species. (A-F) Scatterplots of log_2_FIC_50_ values for all 28 drug pairs tested against (A, B) *Pa* PaO1, (C, D) *Ab* ATCC17978 and (E, F) 45 drug pairs tested against *Kp* ATCC43816. X-values represent log_2_FIC_50_ in CAMHB, while y-values represent log_2_FIC_50_ value in nutrient-depleted media, (A, C, E) M9Glu and (B, D, F) UMM. Lines parallel to the x-axis and y-axis indicate the boundaries of additivity (log_2_FIC_50_ from −0.19 to 0.26, see Materials and Methods). Combinations that fall in the upper-left and lower-right sections of each graph indicate discordant interactions between results in CAMHB and results in the nutrient-depleted medium (marked with a D in the key on the left). Combinations that fall in the gray shaded regions are synergistic in nutrient-depleted media but additive or antagonistic in CAMHB; combinations for which this occurs in one or more species are bold-faced and underlined in the list of combinations on the left.

### Specific antibiotics were associated with synergistic interactions in nutrient-depleted media

We next evaluated if specific antibiotics were more likely to be impacted by changes in media and if single drugs were responsible for higher rates of synergistic interactions dependent on growth medium. To investigate this, we first determined which combinations showed significant differences in log_2_FIC_50_ scores in different media conditions with the same species. The results are shown in Fig. 3 with statistically significant differences between combinations indicated with a teardrop.

**Figure 3:**
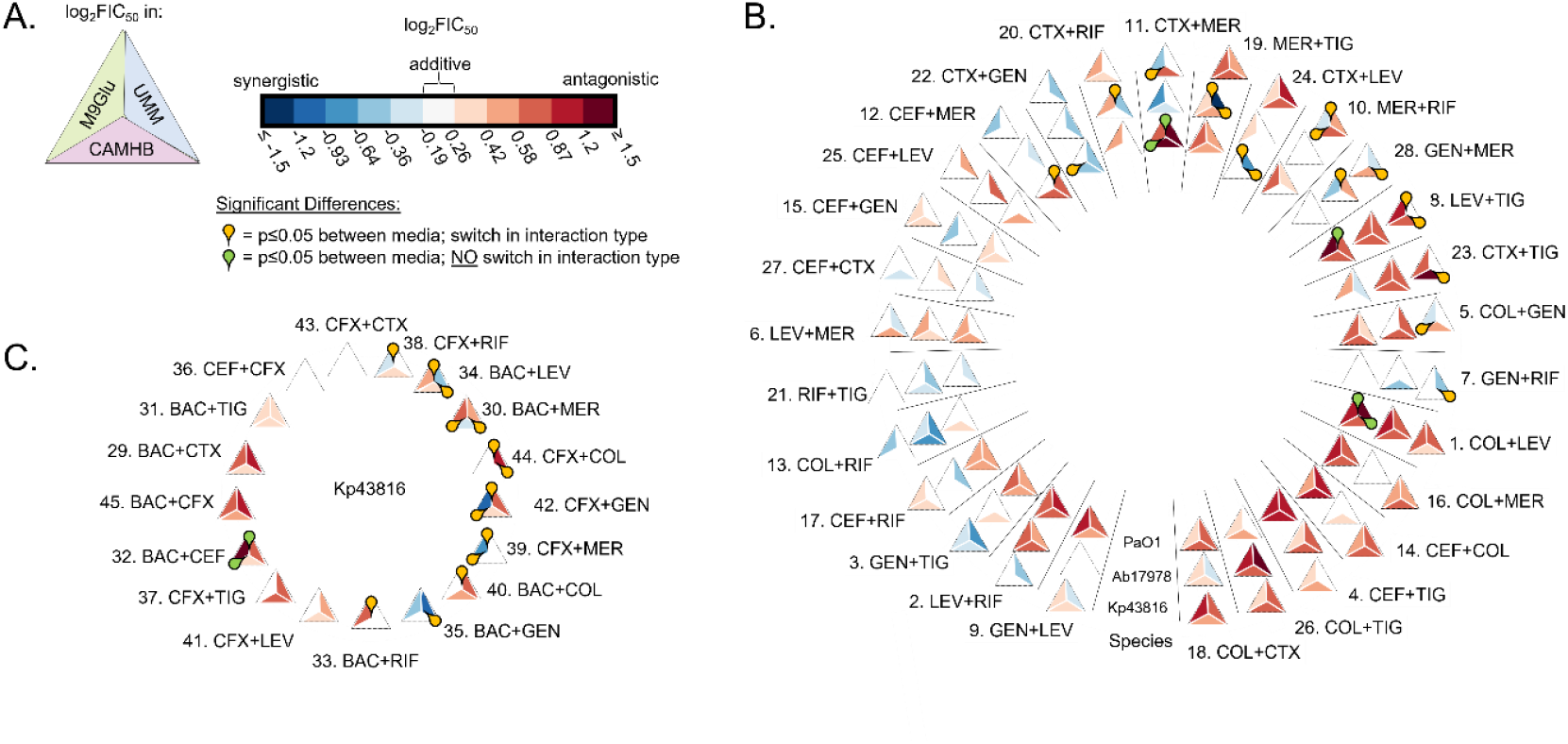
Different species show different degrees of variation across media conditions. (A) Triangle diagram represents how the log_2_FIC_50_ data is depicted in (B) and (C) with UMM value on the top right, M9glu value on top left and CAMHB on the bottom. Log2FIC50 values are reported as in Figure 1. (B, C) Yellow teardrops indicate significant differences (p≤0.05) between media where combinations change interaction type (ex. switch from synergy to antagonism between media); green teardrops indicate significance for combinations that do not change interaction type. Significance was based on a 2-way ANOVA using Tukey’s multiple comparison post-test (α = 0.05), using the log_2_FIC_50_. (B) The outer ring of triangles represent log_2_FIC_50_ data of combinations tested in Kp, the middle ring represents combinations tested in Ab, and the inner ring represents the combinations tested in Pa. (C) The log_2_FIC_50_ data for combinations only tested against Kp.

First, we considered the 28 drug pairs that we tested in all three species (Fig. 3B). For Ab (Fig. 3B middle ring), there were 4 instances of significant differences between interaction measurements, and in all 4 cases the type of interaction (synergy, additivity, antagonism) for a combination switched between two media (yellow teardrops). For Pa (Fig. 3B, inner ring), there were nine instances of significant differences between interaction measurements in two media, but in five of those cases the interaction type did not change between the two media (green teardrops). Finally, for Kp (Fig. 3B, outer ring), there were nine instances of significant differences, and in all cases the interaction type switched (yellow teardrops). Among the additional combinations tested in Kp (Fig. 3C), we saw sixteen significant differences, and the interaction type changed for fourteen of those cases (yellow teardrops) and stayed antagonistic for two cases (green teardrops). Thus, significantly different interaction type switches between media were more frequent in Kp than Ab or Pa, mirroring the same trend observed with the Pearson correlation coefficients where there was the poorest correlation for Kp when comparing impact of antibiotics between different media types (Fig. 1E).

Next, we evaluated whether some drugs were over-represented among significantly different combinations that had instances of switching interaction type among media (e.g., additive to antagonistic or synergistic to antagonistic, Fig. 3, yellow teardrops). Because two drugs in the dataset (trimethoprim-sulfamethoxazole and cefixime) were tested only against Kp, we converted the actual number of interaction switches to a percentage of the total possible interaction switches between media types, for combinations containing that drug. The total number of possible interaction switches was 27 for trimethoprim-sulfamethoxazole and cefixime and 69 for the other eight drugs (see Materials and Methods). The results for all ten drugs are shown in Fig. 4A (yellow bars). Combinations involving trimethoprim-sulfamethoxazole, cefixime, meropenem, and gentamicin were more likely to show a significant difference in interaction type switch between media. We also calculated what subset of the instances of switching involved synergy - i.e., they were not a switch from additivity to antagonism (Fig. 4A, black bars). Of these, over 80 percent of the switches with gentamicin and meropenem involved a change to or from synergy (little differences between black bars and yellow bars, Fig. 4A).

**Figure 4:**
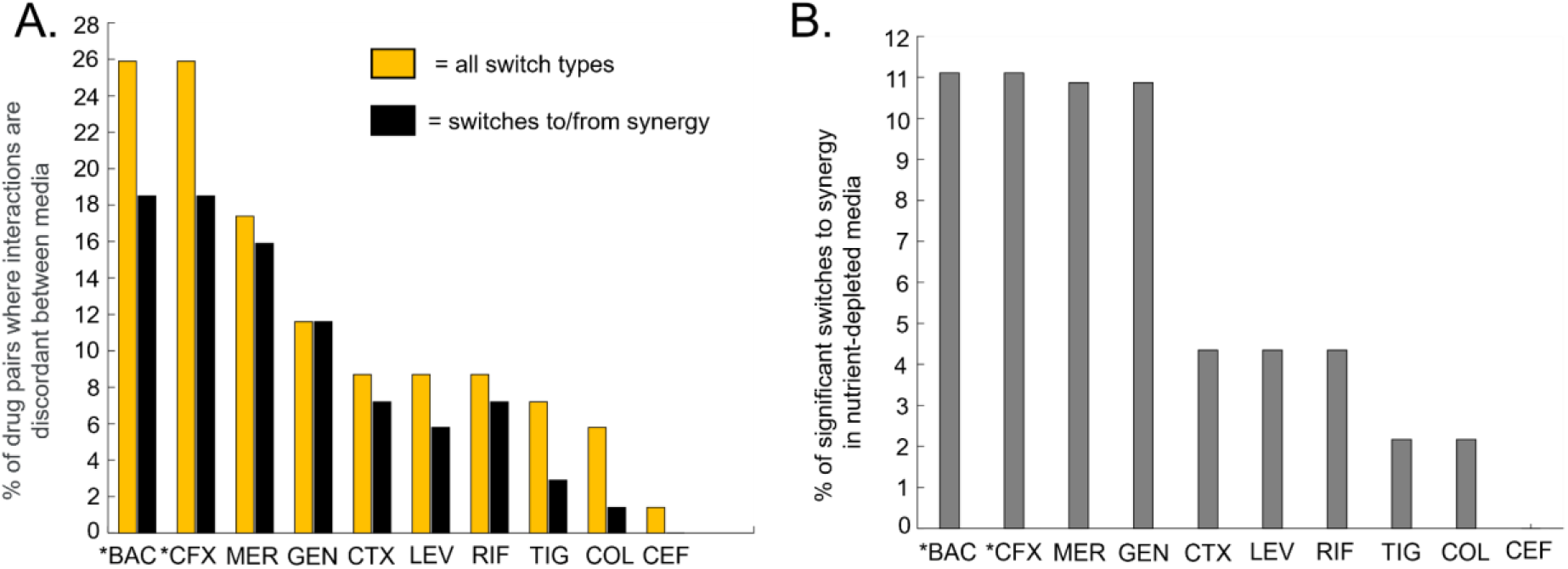
Some drugs were more frequently observed in combinations that show a significant difference in interaction between media. An asterisk indicates a drug that was only tested against Kp. (A) Yellow bars: the percentage of combinations involving each drug that showed a statistically significant log_2_FIC_50_ interaction type switch (ex. synergistic to antagonistic) in different media conditions (yellow bars). Black bars: the percentage of combinations involving each drug that showed a statistically significant log_2_FIC_50_ interaction type switch to or from synergy. (B) The percentage of combinations involving each drug that switched from additivity or antagonism in CAMHB to synergy in nutrient-depleted media (M9Glu or UMM).

We determined the subset of instances that involved switching from additivity or antagonism in CAMHB to synergy in a nutrient-depleted media for each antibiotic (Fig. 3B and 3C, yellow teardrops). To do so, the percentage of switches to synergy by dividing by the total number of potential switches between CAMHB and M9Glu and between CAMHB and UMM was calculated. Trimethoprim-sulfamethoxazole and cefixime had the highest percentage of significant switches to synergy in a nutrient-depleted media (Fig. 4B), with the caveat that they were only tested in Kp. Of the eight drugs tested in all three species, meropenem and gentamicin had the highest percentage of significant switches to synergy in nutrient-depleted media (five switches for meropenem and five for gentamicin). Thus, when testing combinations involving meropenem, gentamicin, as well as trimethoprim-sulfamethoxazole and cefixime in Kp, nutrient-depleted media revealed synergies not observed in CAMHB.

Our *in vitro* data indicate that growth conditions influence the likelihood of observing synergy in combinations. The dependency on growth condition was stronger in Ab and Kp (Figs. 2, 3, 4A). Furthermore, certain antibiotics were more likely to impact medium-dependent synergies. This raises the question of whether *in vitro* data from specific media better reflect *in vivo* outcomes of drug combinations for specific infection types.

### Drug interactions in M9 glucose medium correlated with *in vivo* outcomes for Ab and Kp

For some species, *in vitro* media conditions had varied influence on drug interaction. For example, there was a strong correlation for Ab between M9Glu and CAMHB whereas there was a poor correlation for Kp between these two media (Figs 1B, C and E). Therefore, we postulated that certain media conditions may better reflect *in vivo* drug interactions or efficacy for Kp. To determine whether measurements of drug interactions in specific media are more predictive of *in vivo* outcomes, we investigated whether drug interactions in Ab or Kp grown in CAMHB or M9Glu better replicated *in vivo* observations of drug combinations in lung infections. We analyzed a set of studies in which drug combinations were tested against Ab or Kp lung infections in mice or rats (Tables 2-3, first column) and interpreted the *in vivo* results using the following criteria and strategies (Tables 2-3, second column). We only evaluated animal studies where data was shown for each antibiotic used alone and in combination, and where CFU was measured from lung tissue (Tables 2-3, column 1). If the combination reduced the bacterial burden substantially more than both the single antibiotics used in monotherapy, we interpreted the result as ‘synergistic’ (Tables 2-3, column 2). Alternatively, if CFUs were similar or worse, we called the combination ‘not more effective’ or ‘less effective’. In some cases, one antibiotic was ineffective against Ab or Kp while a second antibiotic was effective. When the ineffective antibiotic further reduced the killing by the effective antibiotic, we considered this ‘potentiating’ and analogous to synergy for this comparison (Tables 2-3, column 2). In several cases, the antibiotic combinations used in the animal studies were not the same as ones used in our dataset (Tables 2-3, columns 1, 3). In these cases, we compared antibiotics within the same class as those in our *in vitro* analyses (Table 1). For example, Zhang et al, use a combination of polymyxin B + meropenem, which we compared to our *in vitro* results with colistin + meropenem (#16).

**Table 2.** Annotations from *in vivo* lung studies of antibiotic combinations used in Ab infections and comparisons to in vitro results

| Source(s), combination, animal model<sup>1</sup> | Interpretation of *in vivo* interaction<sup>2</sup> | Combination (#) from our dataset<sup>3</sup> | Observed *in vitro* interaction for Ab17978 in |  |
| --- | --- | --- | --- | --- |
|  |  |  | CAMHB | M9Glu |
| Fan et al. 2016<br>(colistin + meropenem, mouse)<br>Zhang et al. 2021<br>(polymyxin B + meropenem, mouse) | synergistic or potentiating | COL+MER (#16) | additive | additive |
| Pantopoulou et al. 2007<br>(colistin + rifampicin, rat)<br>Fan et al. 2016<br>(colistin + rifampicin, mouse)<br>Song et al. 2009<br>(colistin + rifampicin, mouse)<br>Zhang et al. 2021<br>(polymyxin B + rifampicin, mouse) | synergistic or potentiating | COL+RIF (#13) | synergistic | additive |
| Montero et al. 2004<br>(imipenem + tobramycin, mouse) | synergistic or potentiating | GEN+MER (#28) | antagonistic | synergistic |
| Sun et al. 2014<br>(meropenem + rifampicin, mouse)<br>Montero et al. 2004<br>(imipenem + rifampicin, mouse)<br>Song et al. 2009<br>(imipenem + rifampicin, mouse) | synergistic or potentiating | MER+RIF (#10) | additive | additive |
| Yilmaz et al. 2012<br>(colistin + tigecycline, rat)<br>Fan et al. 2016<br>(colistin + tigecycline, mouse) | not more effective together | COL+TIG (#26) | antagonistic | antagonistic |
| Joly-Guillou et al. 2000<br>(amikacin + levofloxacin, mouse) | less effective in combination | GEN+LEV (#9) | additive | antagonistic |
| Montero et al. 2004<br>(rifampicin + tobramycin, mouse)<br>Song et al. 2009<br>(rifampicin + amikacin, mouse) | less effective in combination | GEN+RIF (#7) | synergistic | additive |
| Joly-Guillou et al. 2000<br>(amikacin + levofloxacin, mouse) | less effective in combination | LEV+MER (#6) | antagonistic | antagonistic |

1. Studies using combinations of antibiotics in a mouse lung infection model or thigh model and CFU in lung were evaluated (58-65).
2. Our interpretation of the interaction of the combination based on the data in these studies. Classified as either synergistic when the combination is more effective than either monotherapy, or if the combination did not significantly reduce lung bacterial burden it is listed as being not more effective than monotherapies. In most animal studies, individual antibiotic doses were combined to make the combination therapy resulting in higher total drug levels for the combination therapies compared to the monotherapies. Thus, increased killing could be considered additive or synergistic.
3. The combination in our data set (Fig. 2) that is being compared to for similar drug class as the ones used in the study

**Table 3.**
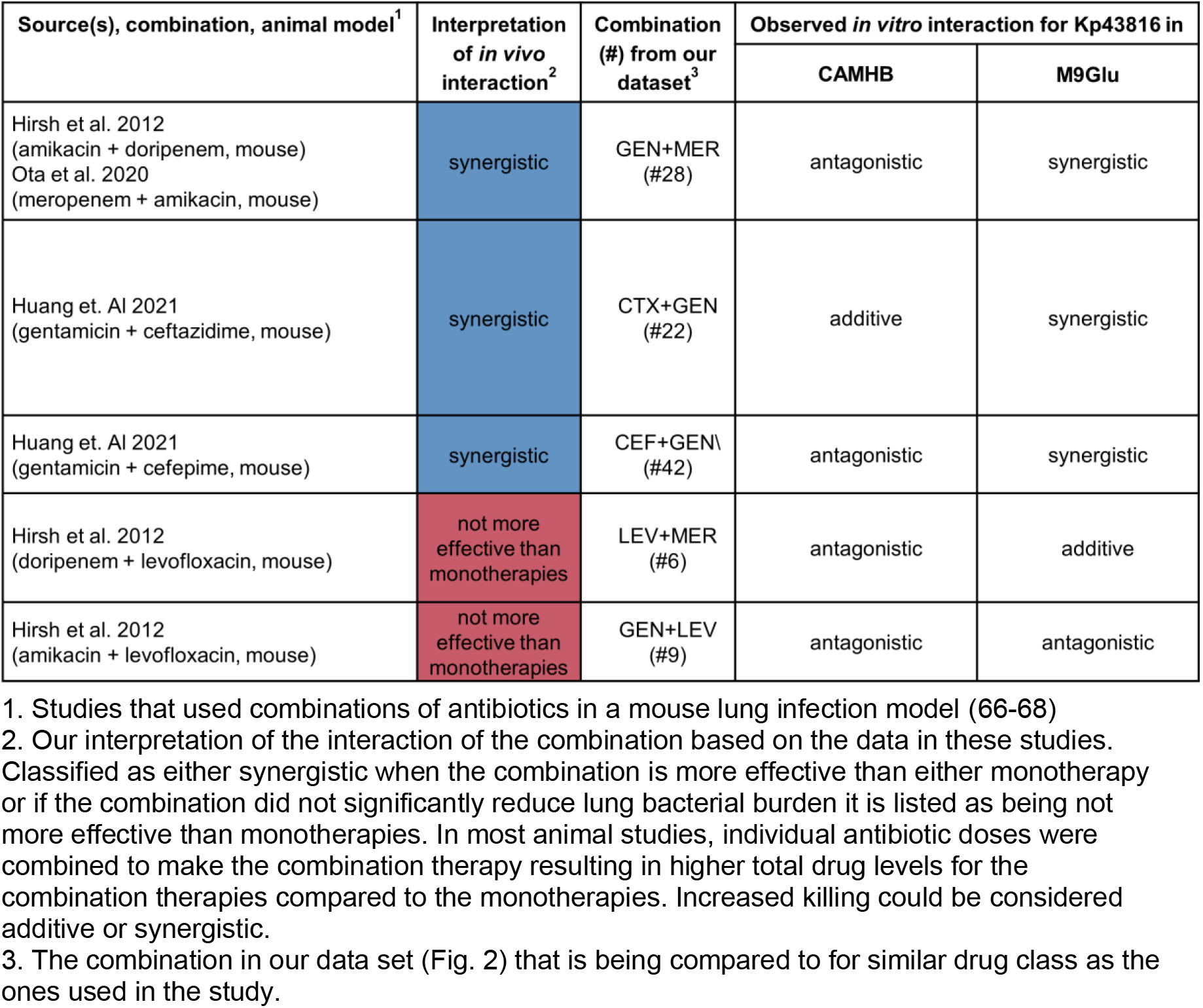
Combinations that were synergistic or antagonistic in mouse *Klebsiella pneumoniae* lung infections

To evaluate whether *in vitro* data in CAMHB or M9Glu better predicts *in vivo* outcomes, we compared the outcome categories from the animal studies with our *in vitro* measures (Tables 2-3, column 5 derived from data from Figs. 2-3). We found that for Kp, M9Glu correlated with all *in vivo* interpretations whereas *in vitro* data from CAMHB would have only predicted one of the combinations (Table 3). By contrast, in Ab, both CAMHB and M9Glu predicted *in vivo* outcomes at similar frequencies (Table 2). Collectively, these results are consistent with the poor Pearson co-efficient comparison for Kp and the high co-efficient for Ab between these media (Fig 1E).

To account for the possibility that drug interactions may vary depending on specific clinical isolates and their individual drug susceptibility patterns, three Ab clinical isolates with a range of resistance profiles (Ab5075, EGA355, and EGA368) were grown in CAMHB and M9Glu and tested against 8 antibiotics combinations (Fig. 5A, B). Ab5075 was highly resistant to gentamicin and meropenem, while EGA355 was highly resistant to levofloxacin resulting in unobtainable IC50 values for these drugs. Thus, for these drugs we tested for potentiation in the relevant strain by adding a constant amount of the resistant drug along with increasing amounts of the sensitive drug and measuring shifts in IC_50_ of the sensitive drug. This shift was reported as a log2 fold-change in IC_50_, with negative log_2_Fold_50_ scores indicating that addition of the resistant drug lowered the IC_50_ of the sensitive drug, despite the resistant drug showing no growth inhibition on its own (Fig 5A, B).

**Figure 5:**
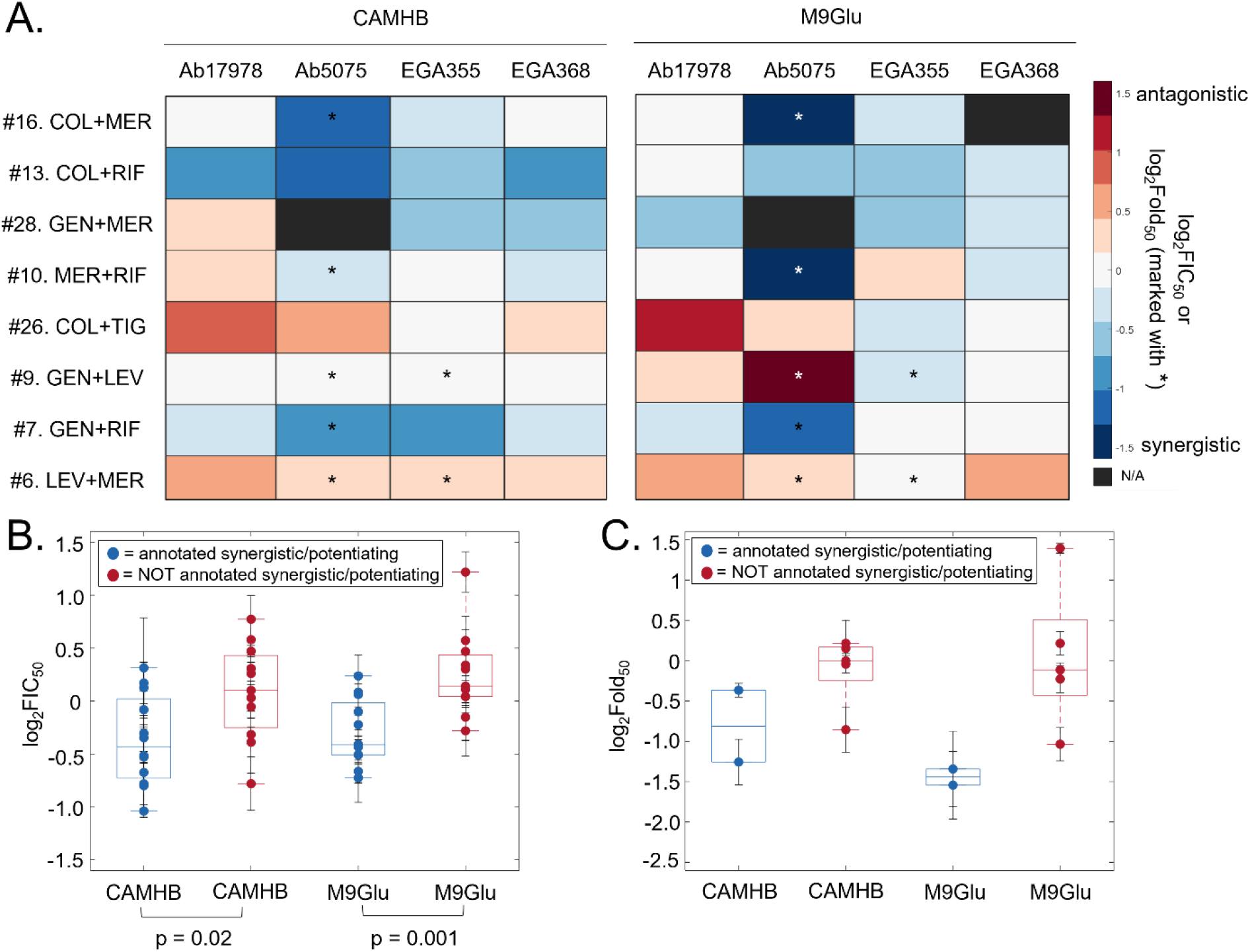
In Ab clinical isolates and lab strain (Ab17978), drug interactions in M9Glu correlate better than those in CAMHB with in vivo studies. (A) log_2_FIC_50_ and log_2_Fold_50_ values for combinations tested against Ab clinical isolates and lab strain (Ab17978) grown in CAMHB and M9Glu. The log_2_Fold_50_ values are indicated with an asterisk. All values are averages of at least three biological replicates. (B-C) Box-and-whisker plots sorted by whether the combinations were annotated synergistic/potentiation (blue) or not (red) for the (B) log_2_FIC_50_ values in CAMHB and in M9Glu or (C) the log_2_Fold_50_. The log_2_FIC_50_ values and log_2_Fold_50_ values are each shown as mean +/− S.E.M. For both comparisons in (B) a two-sample t-test was used with a significance level of α < 0.05.

To evaluate the possibility that drug interactions may vary depending on specific clinical isolates and their individual drug susceptibility patterns, we expanded the number of Ab strains evaluated and compared their responses *in vitro* to our interpretation of the *in vivo*. Three Ab clinical isolates with a range of resistance profiles (Ab5075, EGA355, and EGA368) grown in CAMHB and M9Glu were tested against nine antibiotics combinations (Fig. 5A). Ab5075 was highly resistant to gentamicin and meropenem, whereas EGA355 was highly resistant to levofloxacin resulting in unobtainable IC_50_ values for these drugs. Thus, for these drugs we tested for potentiation in the relevant strain by adding a constant amount of the resistant drug along with increasing amounts of the sensitive drug and measuring shifts in IC_50_ of the sensitive drug. This shift was reported as a log_2_ fold-change in IC_50_, with negative log_2_Fold_50_ scores indicating that addition of the resistant drug lowered the IC_50_ of the sensitive drug, despite the resistant drug showing no growth inhibition on its own (Fig 5A, blocks marked with asterisks). Drug combinations are listed in same order as Table 2 with the 4 ‘*in vivo* synergistic’ combinations on top. Collectively for all 4 strains, more synergistically combinations were observed in the top 4 drug-pairs (10/15 for CAMHB and 10/14 for M9glu) than in the bottom 4 (4/16 for each medium).

To compare the drug responses of these isolates *in vitro* more quantitatively to our interpretation of the *in vivo*, we grouped combination pairs by whether they were predicted to be synergistic or not *in vivo* and compared the log_2_FIC_50_ scores between M9Glu and CAMHB from all strains (Fig 5B-C). There was a greater difference between the log_2_FIC_50_ scores of combinations that were annotated as synergistic (blue) or not (red) *in vivo* when the combinations were measured in M9Glu (p = 0.001) compared to CAMHB (p=0.02). This suggests that M9Glu may be more predictive than CAMHB for Ab strains, but that CAMHB was also predictive. These results are concurrent with the Pearson coefficient observed between M9Glu and CAMHB for Ab (Fig. 1E, R=0.75, p=4.7 x 10-7) as well as the clustering of drug interactions for Ab in M9Glu and CAMHB (Fig. 1B, subgroup III). “

In summary, for this collection of Ab strains, *in vitro* testing in M9Glu was slightly more predictive of *in vivo* outcomes. However, both M9Glu and CAMHB reflected *in vivo* outcomes even when the isolate being tested was highly resistant to one of the drugs in the combination. By contrast, retrospective analyses for Kp strongly suggest that a test medium of M9Glu more clearly differentiates between combinations that are synergistic or antagonistic *in vivo* against Kp lung infection, compared to CAMHB.

### Drug combination outcomes in a Kp mouse lung infection model were predicted by *in vitro* measurements in M9+glucose but not CAMHB

To further probe the ability of *in vitro* media conditions to predict the efficacy of a drug combination *in vivo,* we adapted a mouse model for Kp lung infection to incorporate antibiotic therapy (20, 21). Though traditional drug therapy is designed with the goal of eliminating bacterial burden, we used subtherapeutic doses of antibiotics with the goal of capturing potential synergies in the treatment of tissue infection by Kp. Specifically, we sought doses of single antibiotics (monotherapy) that reduced the lung bacterial burden significantly compared to a vehicle control, but where the bacterial burden remained at detectable levels. If combination treatments were more effective, we would expect fewer colony forming units (CFUs) recovered versus the single doses.

We tested the hypothesis that M9Glu is better able to predict *in vivo* outcomes by testing two combinations of antibiotics that were synergistic in M9Glu, cefixime + meropenem (#39) and cefixime + gentamicin (#42), but additive or antagonistic, respectively, in CAMHB. These combinations were chosen because they were statistically significant in vitro (Fig. 3). Initial testing was done to identify roughly equipotent doses that met the criteria for subtherapeutic doses. Doses of 10mg/kg of meropenem, 5mg/kg of cefixime, and 2 mg/kg of gentamicin given at 14 hours post-infection resulted in a lung bacterial burden between 10^5^ and 10^6^ CFUs 22 hours post-infection after intranasal inoculation of 10,000 CFU whereas untreated controls ranged between 10^7^-10^8^ CFUs. This bacterial burden in treated mice was significantly lower than the non-treated vehicle control while still being 2-3 logs higher than the limit of detection for this assay (Fig. 6).

**Figure 6.**
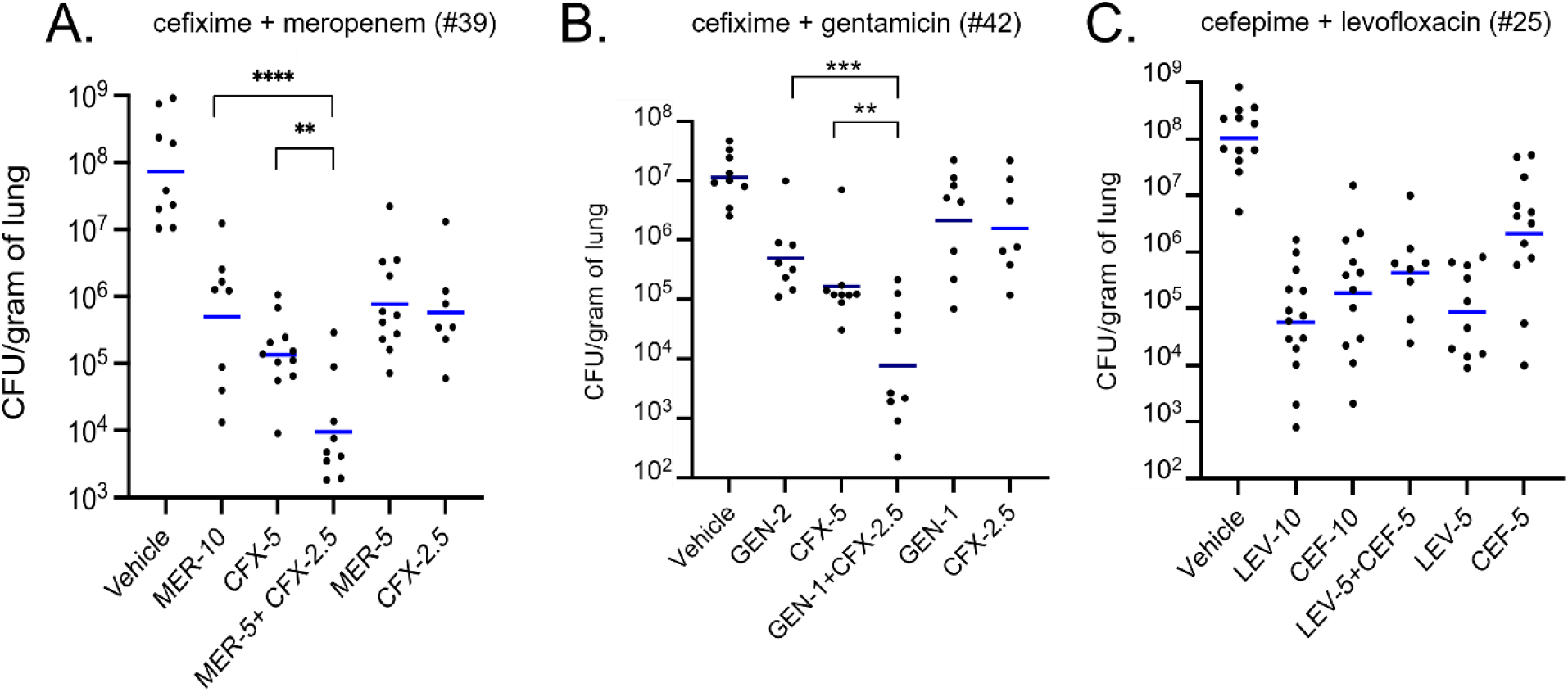
Drug combinations identified as synergistic in M9+Glucose, but not CAMHB, significantly reduce lung bacterial burden during mouse lung infection by *K. pneumoniae.* (A-C) Swiss Webster wild-type mice (black circles) were infected via intranasal route with 10,000 CFUs of Kp43816 and infection was allowed to proceed for 14 hours at which point mice were treated with either DMSO or indicated doses of drugs (in mg/kg) via intraperitoneal injection. Mice receiving meropenem were given a second dose at 18hr due to its short *in vivo* half-life (57). Lungs were harvested after 22 hours post infection and plated for bacterial burden (CFU/gram of lung). Blue lines indicate geometric means. Data for each drug combination group was compiled from n = 3 independent experiments with 3-4 mice in each group. Statistical analysis was done by two-way ANOVA with Bonferroni corrections.

To translate the additivity model used in DiaMOND and compare drug combination therapies to monotherapies *in vivo*, roughly equipotent doses of antibiotics were used for monotherapies and compared to combinations of two drugs each used at half the equipotent dose. For example, 5mg/kg of meropenem + 2.5 mg/kg cefixime was compared to 10 mg/kg meropenem as well as 5 mg/kg cefixime. When either cefixime + meropenem (#39) or cefixime + gentamicin (#42) was used to treat Kp-infected mice, the combination therapy significantly reduced lung bacterial burden compared to their respective monotherapies (Fig. 6A, B). In addition, the inclusion of the individual components of the antibiotic combination doses on their own allowed for the quantification of drug interaction via a modified Bliss independence score (24) using log_10_ transformed values for CFU (see Materials and Methods). In brief, the Bliss independence model compares the observed effect of the combination to an expected inhibitory effect of the combination which assumes the two drugs act independently; positive scores indicate synergistic interactions while negative Bliss scores indicate antagonistic interactions. Using this log-transformed Bliss independence statistic, synergistic Bliss interactions scores of 0.11±0.02 for cefixime + meropenem (#39) and 0.24±0.07 for cefixime + gentamicin (#42) were calculated. Taken together, the significant reduction in lung bacterial burden by the combinations in addition to the positive Bliss scores indicate that these two combinations were acting synergistically in the mouse lung. Therefore, the drug interactions *in vivo* are more closely correlated with the *in vitro* measurements in M9Glu rather than the additive or antagonistic interactions measured in CAMHB.

To evaluate whether combination therapies broadly acted more effectively than monotherapies in this model regardless of drug interactions measured *in vitro*, cefepime + levofloxacin (#25) was used for *in vivo* testing – a combination that acted additively in both M9Glu and CAMHB. Doses of 10mg/kg for both drugs were chosen for monotherapy and 5 mg/kg of each for combination therapy. The combination of cefepime + levofloxacin (#25) was not significantly different from either monotherapy alone and had an antagonistic log-transformed Bliss interaction score of −0.22± 0.07 (Fig. 6C). Together, these experiments demonstrate that by using subtherapeutic antibiotic doses, the mouse model resolved differences in single versus combination drug therapy. Additionally, these data demonstrate that for both the combinations of cefixime + meropenem (#39) and cefixime + gentamicin (#42) M9Glu medium, an *in vitro* medium more nutritionally restricted similar to the lung environment, was better able to predict *in vivo* behavior when compared to a standard rich media.

## Discussion

Our results demonstrate that drug interactions differ considerably across species. Within species, drug interactions also vary amongst growth conditions. Furthermore, among the panel of drug combinations tested, we observed little to no correlation between three Gram-negative species, in any of the media tested (Fig. 1D) and no drug combination was synergistic across all three species and growth conditions. This discordance in response to drug combinations across different species raises the important and unanswered question of what the best way to assess combination therapies is. Our observation that one cannot extrapolate from one bacterial species to another has also been observed by other investigators. Brochado and colleagues tested pairwise combinations from a broad array of antibiotics classes against *E. coli*, *S. Typhimurium* and Pa grown in Lysogeny Broth, and found that more than 70% of their tested drug interactions were species-specific (25). This variation in response to antibiotic combinations amongst different species could be due to differences in antibiotic uptake (26, 27), cell wall permeability (28), and/or cellular processes when grown in complex nutrient environments (29). Consistent with the latter idea is our observation that differences in drug interactions between species were the least evident in the simplest medium, M9Glu (Fig. 1D). Collectively, our findings indicate that there may not be a “golden” combination that will be synergistic across a range of species and infection sites. Given the impact of species-specific physiology on drug interactions, a more tailored strategy focused on the pathogen and sites of infection may need to be considered.

Among our systematic drug interaction measurements, antagonism was overall more frequent than synergy (Fig. 1B, 1C), which is in agreement with studies of other species (13, 25, 30–33) as well as with cancer therapies (34). However, ceftriaxone + gentamicin was synergistic across all three species in UMM (Fig. 1B). There are other *in vitro* and clinical evidence of synergy for combinations of beta-lactams and aminoglycosides in both Gram-positive and Gram-negative bacteria. For example, synergy was observed in the more rapid clearance of *Staphylococcus aureus* from cardiac vegetations in a rabbit endocarditis model by penicillin combined with gentamicin, compared to either drug alone (35); a similar effect was also observed with *Streptococcus sanguis* in the rabbit endocarditis model (36). Synergy was also observed with amoxicillin in combination with gentamicin when used to treat various strains of *Streptococcus pneumoniae* in a mouse pneumonia model that varied in their penicillin susceptibility (37). In these cases, the cephalosporin is believed to weaken the cell wall allowing better penetration of the aminoglycoside (38–40). Some *in vitro* studies with Pa have shown synergy with a beta-lactam and aminoglycoside (41), but in the case of Pa synergy appears to depend on the strain as well as the specific identity of the beta-lactam with an aminoglycoside in combination (42, 43). Though we did not detect synergistic interactions in every case between gentamicin and the beta-lactams tested, their overall interactions skewed towards additivity/synergy (22 pairs) rather than antagonism (8 pairs). This is in stark contrast to the overall skew towards antagonism in the data set (158/303 possible combinations). These observations further support the idea that these antibiotics may be particularly beneficial for the treatment of complex urinary tract infections.

Our dataset allowed us to take an in-depth look at how drug interactions vary across growth conditions and in different species. Though we focused on statistically significant interaction differences (Fig. 3, 4), we reported all media-to-media interaction differences (Fig. 1) for consideration. Drug interactions may be dependent on media for a variety of reasons, including differences in metabolic state (44, 45), the activity of efflux pumps (46, 47), and stress response pathways which can change depending on media condition (48, 49). For Kp and Ab, combinations that included gentamicin or meropenem were more likely to change to synergistic when moving from a rich medium (CAMHB) to non-rich media (M9Glu or UMM) (Fig. 4B). This highlights the importance of testing combinations involving these drugs in non-rich growth conditions which may better reflect *in vivo* outcomes for some types of Kp and Ab infections. However, this trend with gentamicin and meropenem was not observed in Pa. For Pa, the overall trend toward antagonism has been observed in at least one other study of a broad range of antibiotics (25). The relatively low discordance in drug interaction across media for Pa may be explained by its metabolic adaptability, minimal nutritional requirements, and ability to grow in a variety of different environments (45). These features combined with a wide array of innate resistance mechanisms (50) suggest that *Pseudomonas* can face challenges from multiple antibiotics concurrently, along with environmental stressors. In contrast, Kp undergoes shifts in metabolism upon growth in glucose or other changes in carbon sources (51, 52), and exposure to subinhibitory amounts of meropenem also shifts the metabolism of Kp (53). It would stand to reason that a reverse of this also occurs, that changes in Kp metabolism will exert an effect on drug interaction.

We used two approaches to evaluate which, if any, *in vitro* medium would best predict *in vivo* efficacy. For Ab and Kp, we compared our *in vitro* measurements of drug interactions to results from mouse and rat lung infection studies (Tables 2, 3, Fig. 5). For Ab, we additionally compared our results with ATCC 17978 to a panel of clinical isolates. In the case of Kp, results in M9Glu aligned with clinical findings whereas for Ab, both M9Glu and CAMHB were correlated with *in vivo* results. However, even for Ab, drug interactions in M9Glu tended to better recapitulate *in vivo* findings compared with interactions in CAMHB (Fig. 5B), even though M9glu is not an exact mimetic of lung conditions. For example, lungs contain detectable, albeit low and insufficient, levels of amino acids but there are no amino acids in M9Glu (21). To further explore the predictive power of *in vitro* measurements, we directly tested whether drug interactions in M9Glu or CAMHB better reflected results for a Kp mouse lung infection model. By using drugs at subtherapeutic levels, we had the resolution to detect enhanced clearance in lung bacterial burden when drugs were used in combination compared to as a monotherapy. This model permitted us to experimentally confirm that M9Glu better predicted drug combinations than CAMHB. Collectively, these analyses indicate that for Kp, M9Glu is better able to predict *in vivo* outcomes in the lungs when compared to CAMHB (or UMM). This further implies that Kp is using a glycolytic program during its growth in the lungs and that these drug combinations are more effective under these conditions. Additionally, our results for cefixime + meropenem (#39) are in accord with previous clinical trial results, providing further evidence for the efficacy of double beta lactam therapy for multi-drug resistant Kp (54).

Traditional drug therapy in mice is often designed with the goal of eliminating the bacterial burden utilizing full doses of each drug together. Our dosing strategy, which uses combinations with half the dose of the monotherapy, was designed to measure *in vivo* drug interactions relative to additivity as a null model (55, 56). This allowed for a more direct comparison between a combination and its respective monotherapies. A potential strength of using subtherapeutic concentrations is the resolution to detect both decreases and increases in bacterial burden when giving a combination of drugs. Although we weighted our drug doses to detect further decreases in bacterial burden when using combinations, this model can be optimized to better capture antagonistic interactions by altering both the doses. Additionally, this dosing strategy using subtherapeutic concentrations can be adapted to test whether other infection site-specific mimetic media can achieve the same recapitulation observed here. For example, do results in UMM better recapitulate interactions in a Kp mouse model for cystitis compared to results in CAMHB? Future work will explore if this is the case. If so, then not only could tissue mimetic media be used to better predict *in vivo* outcomes in corresponding infection sites, but results of a panel of tissue mimetic media could be used to identify combinations that perform well across multiple sites in more complex infections.

Together, our findings have several implications. First, it should not be assumed that a drug combination will behave the same way in different growth conditions. However, for some species such as Pa, testing different media conditions may not be necessary, while for other species such as Kp infection sites may need to be carefully considered when choosing an appropriate combination therapy. For multi-site infections, choosing a combination that performs well across a range of growth conditions might be the best strategy. In addition, there is no consistent pattern of media-to-media variation between species; for Kp and Pa, responses in CAMHB and UMM were more similar, and for Ab, responses in CAMHB and M9Glu were more closely related (Fig. 1E). Thus, even for species like Ab and Kp for which changes in media affect drug combination response, there is variation between species in the magnitude of the effect that a specific media will have on drug responses. Our study suggests that informed use of combination therapies should take account of species and infection sites, and furthermore, that for some species growth conditions may have an outsized effect on combination interactions, as we have started to observe with this work. More studies are needed to further characterize the effect of species and growth conditions on drug interactions, to inform the design of better combination therapy.

## Materials and Methods

### Strains, antibiotics, and growth conditions

Strains used in this paper include Ab ATCC 17978 (a generous gift from the lab of Ralph Isberg at Tufts University), Pa PaO1 (a generous gift from the lab of Paul Blainey at the Broad Institute), and Kp ATCC 43816, as well as three Ab clinical isolates. Ab5075 is a well-characterized, extensively drug resistant (XDR) isolate from a Walter Reed Army Medical Center patient between 2008 and 2009 (Jacobs et al 2014, Thompson et al 2014, Zurawski et al 2012). Susceptibility and resistance information for this strain was obtained from Wu et al. (2015) and Jacobs et al. (2014). EGA355 and EGA368 (obtained from Eddie Geisinger in the lab of Ralph Isberg at Tufts University) are two Ab strains that were isolated from patient sputum samples in 2013 and 2014, respectively, by the Tufts Medical Center Microbiology Laboratory. Species confirmation and MLST strain type (ST2) were determined by whole-genome sequencing.

Ten antibiotics were used in this study. Cefepime, colistin, ceftriaxone, gentamicin, levofloxacin, trimethoprim, sulfamethoxazole, cefixime, and meropenem were obtained from Sigma. Rifampicin and tigecycline were obtained from T.C.I. Chemicals. For *in vitro* studies trimethoprim and sulfamethoxazole were mixed at a 1:20 ratio. Cation-Adjusted Mueller Hinton II Broth (CAMHB) was purchased from Becton-Dickinson (BBL, Sparks, MD, USA) and prepared according to the manufacturer’s instructions. M9 Minimal Salts 5x was purchased from Becton-Dickinson (Difco, Sparks, MD, USA), and M9 Minimal Medium (M9Glu) was prepared according to the manufacturer’s instructions (including addition of 0.5% glucose). M9 was supplemented with 0.6μM Fe(II)SO_4_ for growing all strains, and with 10mM NaC_2_H_3_O_2_ for growing Ab and Pa strains. Urine mimetic media (UMM) was prepared according to the recipe of Brooks and Keevil (1997) and supplemented with 0.6μM Fe(II)SO_4_ and 0.01% glucose when used for growing Kp ATCC 43816.

### Drug interaction measurement with DiaMOND Assays

First dose centering experiments were performed to determine the IC90 values of each antibiotic for each strain in each medium. The same experimental protocol was used for both DiaMOND and dose centering experiments: a culture was grown overnight to saturation in the medium to be tested at 37°C with shaking, then 6μl of culture was used to inoculate 3ml fresh media, and this day culture was grown at 37°C with shaking until it reached mid-log (OD_600_=0.2-0.5). This day culture was then diluted to OD_600_=0.001, and 50μl culture was added to each of the non-edge wells of 384-well microplates, which had drugs dissolved in DMSO (ceftriaxone, levofloxacin, meropenem, rifampicin, tigecycline, trimethoprim, sulfamethoxazole and cefixime), or 0.1% Triton-X100 in water (cefepime, colistin, and gentamicin), pre-added to the plates using the HP D300E Digital Dispenser. Increasing amounts of single drugs and increasing total amounts of pairs of drugs were used to generate dose response curves for single drugs and pairs of drugs. For each plate, ≥ 4 wells were left untreated (no drug added), and 4-8 wells were treated positive controls, which received 3x MIC of one of the drugs tested. These controls were used for calculating the Z score, see Data Processing and Quality Control below. Then 50μl sterile media was added to each edge well of the 384-well plates. Plates were grown overnight (18-20 hours) with 37°C with shaking. The OD_600_ of each well was measured using a Biotek Synergy HT Microplate Reader. One biological replicate was performed for the dose centering for each species and growth condition, and ≥ 3 biological replicates were performed for each single drug and pairwise combination tested against each strain and medium (Fig. S1 and S2).

### Data processing and quality control

All data analysis was performed in Matlab. The data for each biological replicate was analyzed separately, and log_2_FIC_50_ values and log_2_FIC_90_ for each biological replicate that passed quality control (see below) are reported in Figs.S1 and S2, respectively. Each reported log_2_FIC_50_ value is the arithmetic mean of log_2_FIC_50_ values reported in Fig. S1.

Processing the OD_600_ data by background-subtraction of the median of medium-only edge wells, normalization to the mean of untreated wells in each plate, fitting of the single and pairwise dose response curves with a three-parameter hill function, and calculation of inhibitory concentration (IC) values based on hill curve parameters was performed as described previously (13). Determination of FIC_50_ scores using the IC_50_ value of the drug pair as well as the IC50 values of the component single drugs following the model of Loewe additivity was done as described previously (13). The Ab clinical isolate Ab5075 was highly resistant to gentamicin and meropenem, and the Ab clinical isolate EGA355 was highly resistant to levofloxacin. So, for combinations including gentamicin or meropenem for Ab5075 (gentamicin + meropenem was not tested for Ab5075) and combinations including levofloxacin for EGA355, the drug to which the strain was highly resistant was treated as a sensitizer, and for the combination dose response curve a constant amount of the sensitizer drug was added to an increasing amount of the other drug in the pair. Instead of calculating the FIC50 score for the drug pair, the fold-change between the combination IC50 and the non-sensitizer drug IC50 was calculated as a measure of potentiation, and in data processing instead of normalizing to the mean of the untreated wells, the wells for the combination dose response curve were normalized to wells treated with only the sensitizer drug. We consider potentiation analogous to synergy because both involve the combination of two drugs showing greater efficacy than the sum of the drugs’ individual effects. Otherwise, we categorized the drug interaction as ‘not more effective’ if killing appeared similarly to the single doses or ‘less effective’ if more CFU were recovered in the combination dose compared to one or both single doses.

To ensure accuracy and consistency, all biological replicates included in the dataset had to pass the following series of quality control criteria. For single drug dose response curves, the R2 of the fitted curve (from which we calculated IC values) had to be ≥ 0.9, and the 384-well plate on which the dose response curve was measured had to have a Z score of ≥ 0.4, to ensure sufficient difference between untreated and treated positive control wells requiring consistent growth in the untreated wells and growth inhibition in the positive control wells. The equation we used for Z score calculations is 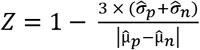 In this equation, 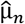 and 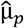 are the average OD_600_ of the untreated and positive control wells, respectively, and 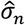 and 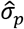 are the standard deviation of the untreated and positive control wells, respectively. We used the same requirements for combination dose response curves for which FIC50 was calculated, with the added criteria that these requirements also had to be met for the component single drugs’ dose response curves, and the angle score for the combination (a measure of how close the single drugs doses were to achieving equipotency) had to be between 23° and 68° (no more than 22° degrees away from 45°, indicating equipotency and exact measurement along the diagonal).

### Determination of additivity range, synergy and potentiation

To experimentally determine the window of additivity in our assays, the range of log_2_FIC_50_ scores obtained by measuring 3-5 drugs from the panel individually in the DiaMOND format with themselves (e.g., a mock combination experiment) against Ab17978, PaO1 and Kp43816 each grown in CAMHB and in M9Glu. For each species in each medium, at least two biological replicate measurements were performed for each drug tested with itself, and the resulting log_2_FIC_50_ scores were used to calculate a 95% confidence interval for additivity for each species in each media. All six of these 95% confidence interval ranges (three species in two media) were within the range of log_2_FIC_50_ = 0.26 and log_2_FIC_50_ = −0.19. Thus, log_2_FIC_50_ scores between −0.19 and 0.26 were considered additive, while scores less than that were considered synergistic and scores greater than that were considered antagonistic.

### Statistical analysis

For each species, we identified the combinations with statistically significant differences in interaction type between growth conditions by performing a 2-way ANOVA with multiple comparisons using Tukey’s multiple comparison post-test (α = 0.05), with the log_2_FIC_50_ scores from all combinations in CAMHB, M9Glu and UMM. Combinations were considered statistically significant if p≤0.05 in log_2_FIC_50_ between two growth conditions.

For each of the 10 drugs tested, we counted the total number of combinations involving that drug that switched interaction type (ex. synergy to antagonism) between two growth conditions, across all the growth conditions and species tested. For comparisons between drugs (Fig. 4), we converted each total to a percentage of all the possible switches in interaction type between growth conditions, across all growth conditions and species. Since trimethoprim-sulfamethoxazole and cefixime were only tested in Kp, there are 27 possible switches for each of these two drugs: 1 species x 3 possible media-to-media comparisons x 9 combinations. For the other 8 drugs, there are 69 possible switches: 2 species (Ab, Pa) x 3 possible media-to-media comparisons x 7 combinations, plus 1 species (Kp) x 3 possible media-to-media comparisons x 9 combinations (since any of these other eight drugs was also tested with trimethoprim-sulfamethoxazole and cefixime in Kp).

### Mouse infections

For infections, 8-12 week old female or male Swiss Webster mice (Taconic) were anesthetized with isoflurane and infected via the intranasal route with 50ul containing 10,000 CFU of stationary phase Kp (ATCC43816) grown overnight in L broth and diluted in sterile PBS (20). Prior to infection, mice were weighed to ensure accurate doses of antibiotic(s). Infection was allowed to proceed for 14 hours. At which point, stated concentrations of antibiotics diluted in 100μl of DMSO were administered via intraperitoneal injection. For combination doses, antibiotics were mixed in 100 μl DMSO. A cohort of mice were given 100μl of DMSO at 14 hours post infection. (Antibiotic concentrations used were based on preliminary experiments (not shown) that identified antibiotic concentrations that reduced bacterial burden 50-500 fold compared to vehicle). Due to the short half-life of meropenem, a 2^nd^ dose was given at 18 hours post infection. All other antibiotics have longer half-lives in mice (57). Mice were euthanized at 22 hours post infection. Lungs were collected, weighed, and homogenized. Homogenates were diluted, plated on L agar plates, and grown at 37°C overnight. CFUs were counted and used to calculate lung bacterial burden per gram of lung. A 2-way ANOVA with Bonferroni’s multiple comparison corrections (α = 0.05) was done on log_10_ transformed data to determine statistical significance using GraphPad Prism. All infections were done at least three times with groups of 2-4 mice/condition and data compiled. To calculate Bliss interaction scores, log_10_ CFU/gram of lung was used to calculate the relative inhibition for each treatment group. These values allowed for the implementation of the Bliss independence model to calculate the expected inhibition if there was no interaction between the two drugs being used (24). To calculate the expected inhibition Eq1 was used, where y_A_ and y_B_ is the observed fractional growth inhibition by drug A and drug B respectively at ½ the dose used for the combination therapy (for example 2.5mg/kg of cefixime and 5mg/kg of meropenem), y_B_ is the observed growth inhibition by drug B. Fractional growth inhibition was calculated by log_10_ transforming the geometric means of the CFU/g of lung for each group of mice and dividing the treated groups by the untreated group. The expected growth inhibition is subtracted from the observed growth inhibition to calculate the Bliss score for the combination.

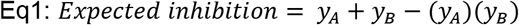

## Acknowledgments

We thank Nhi Van, Jonah Larkins-Ford, Yonatan Degefu, and Talia Greenstein for coding assistance and experimental advice. We thank Pathricia Leus and Rachel Ende for critical reading of manuscript. YM was supported by 4T32AI007422 award to Ralph Isberg from NIH (NIH NIAID); ALM was supported by NIGMS grant 1K12GM133314 awarded to Dr. Claire Moore. KPD, JM and BA were supported by NIH NIAID U19142780 awarded to Deborah T. Hung. JM was also supported by NIH NIAID AI169786.

## Supporting Information for

**Figure S1:**
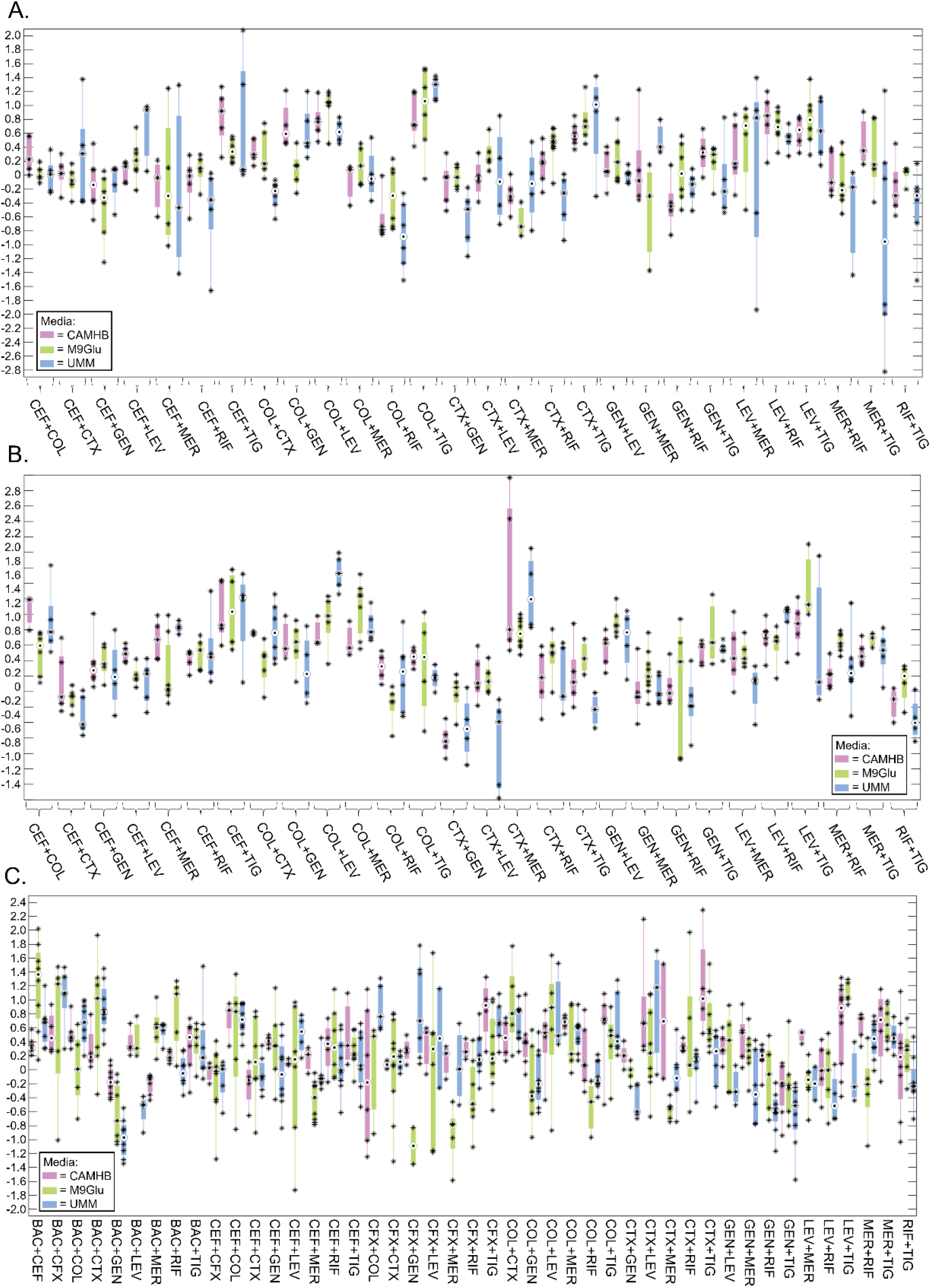
Biological replicates of pairwise drug combination log_2_FIC_50_ measurements against (A) *Acinetobacter baumannii* ATCC17978, (B) *Pseudomonas aeruginosa* PaO1, and (C) *Klebsiella pneumoniae* ATCC43816, each grown in CAMHB (purple), M9Glu (green), and UMM (blue). Box plots depict the median (central circle), 25^th^ and 75^th^ percentiles (edges), and whiskers extend to the largest and smallest replicate values. Individual replicate values are marked with a black asterisk.

**Figure S2:**
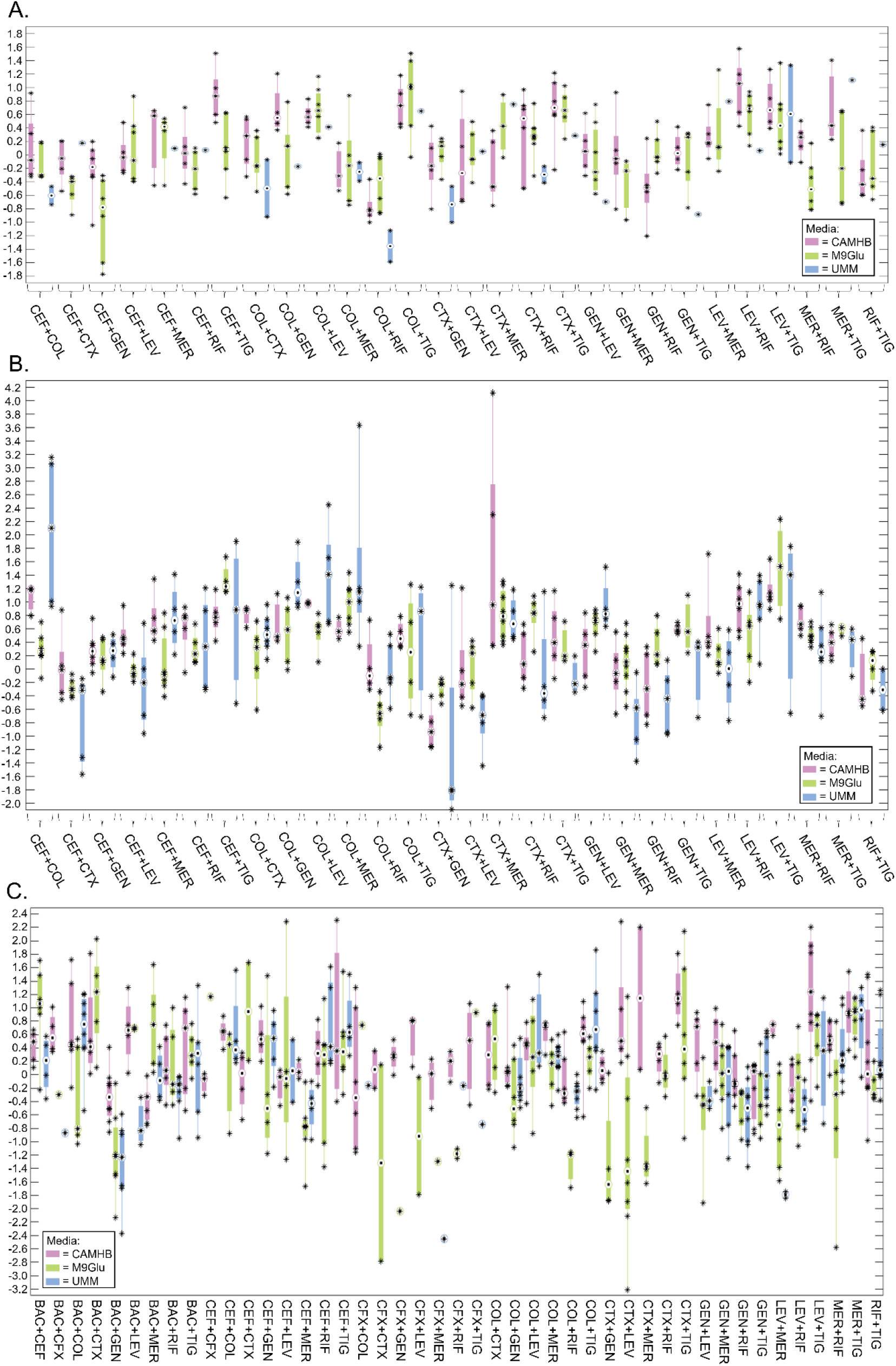
Biological replicates of drug combination log_2_FIC_90_ measurements against (A) *Acinetobacter baumannii* ATCC17978, (B) *Pseudomonas aeruginosa* PaO1, and (C) *Klebsiella pneumoniae* ATCC43816, each grown in CAMHB (purple), M9Glu (green), and UMM (blue). Box plots depict the median (central circle), 25^th^ and 75^th^ percentiles (edges), and whiskers extend to the largest and smallest replicate values. Individual replicate values are marked with a black asterisk.

